# PathExt: a general framework for path-based mining of omics-integrated biological networks

**DOI:** 10.1101/2020.01.21.913418

**Authors:** Narmada Sambaturu, Vaidehi Pusadkar, Sridhar Hannenhalli, Nagasuma Chandra

## Abstract

**Motivation:** Large scale transcriptomic data are routinely used to prioritize genes underlying specific phenotypes. Current approaches largely focus on differentially expressed genes (DEGs), despite the recognition that phenotypes emerge via a network of interactions between genes and proteins, many of which may not be differentially expressed. Furthermore, many practical applications lack sufficient samples or an appropriate control to robustly identify statistically significant DEGs.

**Results:** We provide a computational tool - PathExt, which, in contrast to differential genes, identifies differentially active paths when a control is available, and most active paths otherwise, in an omics-integrated biological network. The sub-network comprising such paths, referred to as the TopNet, captures the most relevant genes and processes underlying the specific biological context. The TopNet forms a well-connected graph, reflecting the tight orchestration in biological systems. Two key advantages of PathExt are (i) it can extract characteristic genes and pathways even when only a single sample is available, and (ii) it can be used to study a system even in the absence of an appropriate control. We demonstrate the utility of PathExt via two diverse sets of case studies, to characterize (a) Mycobacterium tuberculosis (M.tb) response upon exposure to 18 antibacterial drugs where only one transcriptomic sample is available for each exposure; and (b) tissue-relevant genes and processes using transcriptomic data from GTEx (Genotype-Tissue Expression) for 39 human tissues. Overall, PathExt is a general tool for prioritizing context-relevant genes in any omics-integrated biological network for any condition(s) of interest, even with a single sample or in the absence of appropriate controls.

**Availability:** The source code for PathExt is available at https://github.com/NarmadaSambaturu/PathExt.

## 1 Introduction

Whole-genome transcriptomic data are routinely harnessed to probe genes and processes underlying specific biological contexts, including diseases (Blumenberg (2019); Jiang *et al.* (2015)). Extracting biological insights from such high-dimensional data remains an important challenge (Esteve-Codina (2018)). A standard approach to interpreting such data is to first identify differentially expressed genes (DEGs) and then to identify enriched functions among such genes (Esteve-Codina (2018)). However, biological phenotypes emerge from complex interactions among numerous biomolecules, resulting in a highly heterogeneous transcriptional landscape, thus adversely affecting the power to detect critical genes and pathways based on DEGs alone. Moreover, such high-coverage data encodes a vast amount of information beyond DEGs, warranting exploration using multiple complementary approaches. Genome-wide molecular interaction networks constructed from experimentally identified physical, regulatory, signaling, and metabolic interactions have shown great promise as a framework for integrating and interpreting such data (Sambarey *et al.* (2017a,b)). The identification of sub-networks in such biological networks, which encode the processes perturbed by a stimulus, or active processes in general, can lead to mechanistic insights, as well as help prioritize genes for intervention (Mitra *et al.* (2013)). Several methods have been proposed to integrate transcriptomic data with biological networks, that identify ‘active modules’ or connected sub-networks which show changes across conditions (Mitra *et al.* (2013)). Despite the availability of interaction data, these methods largely rely on network scoring schemes which prioritize DEGs (Mitra *et al.* (2013)). However, in many practical scenarios including clinical settings, lack of appropriate controls or sufficiently large number of samples preclude robust identification of statistically significant DEGs (Stretch *et al.* (2013)).

In this work, to complement the conventional differential expression-based analyses, we provide PathExt, a path-based approach to mining omics-integrated biological networks. PathExt uses a network weighting scheme that prioritizes edges/interactions rather than nodes/genes, and identifies differentially active paths when comparing conditions, or highly active paths when studying a single condition. The sub-network comprised of these differential paths, referred to as the TopNet, captures the genes and pathways characterizing the biological condition under study. Deviating from traditional approaches to active sub-network identification, PathExt does not use the selection of a connected module as a constraint. Rather, the method results in a well-connected sub-network, reflective of the interconnectedness of biological processes responding to any stimulus.

PathExt can be used to address the following biologically important questions: (i) What are the most significantly differential paths between conditions, and what are the most critical genes underlying the differentially active paths (note that the critical genes themselves may not be differentially active)?; (ii) What is the response to a given perturbation?; and (iii) What are the most active paths and processes in a condition for which there is no appropriate control?

We demonstrate the wide applicability of PathExt by applying it to two diverse sets of case studies. (a) Exposure of the pathogen Mycobacterium tuberculosis to 18 antibacterial drugs, where only one sample is collected for each such exposure. We find that the TopNet for each sample reveals the pathways known to be affected by the corresponding drug. (b) Transcriptomic data for 39 human tissues. Application of PathExt reveals tissue-relevant genes and processes despite the absence of a clear control. In all applications, we find that the TopNet forms a well-connected graph (not expected by chance). Overall, PathExt is a general framework for the integration and analysis of knowledge-based biological networks and omics data, to reveal context-relevant genes and processes. This can be done even with a single sample, or in the absence of appropriate controls. We provide the open source PathExt tool at https://github.com/NarmadaSambaturu/PathExt.

## 2 Methods

### 2.1 PathExt

We provide an overview of PathExt in Figure 1. The inputs to PathExt are (a) a directed gene network and (b) gene-centric omics data for the conditions of interest. The omics data can represent a variety of quantities pertaining to the node, such as gene expression level, differential expression, protein, metabolite level, etc., in one or more conditions. The output of PathExt is a sub-network, that we refer to as the TopNet, consisting of the most significant differential or active paths, and is interpreted based on the application context.

**Figure 1:**
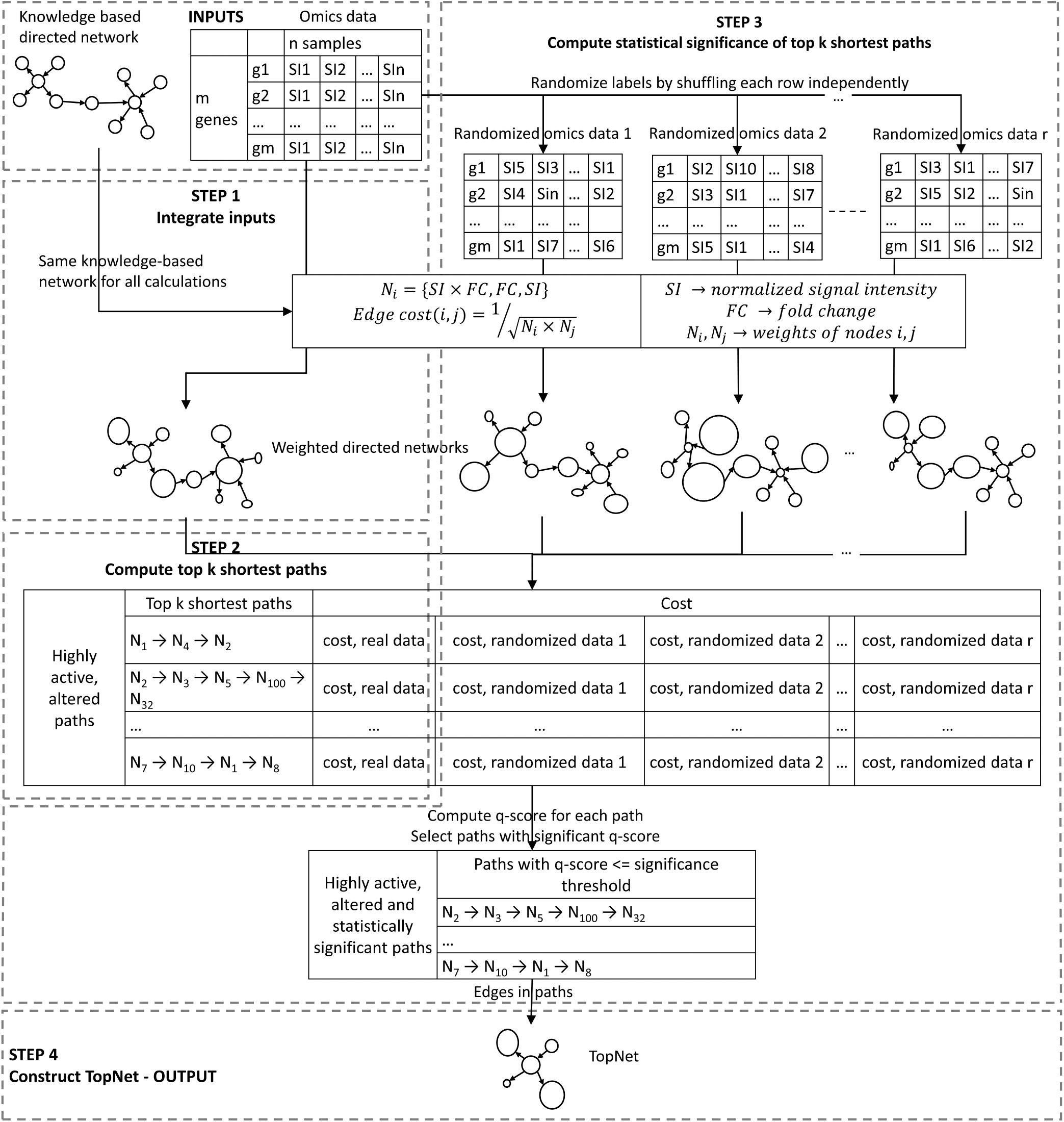
PathExt overview. PathExt uses a knowledge-based directed network and omics data as inputs, and outputs a sub-network consisting of context-relevant genes and processes, referred to as the TopNet.

PathExt can be used to interrogate any combination of knowledge-based networks and omics data. For clarity, we describe the steps for a protein-protein interaction network (PPIN) and gene expression data. The pipeline consists of the following steps (Figure 1): (1) Integrate inputs, (2) Compute top k shortest paths, (3) Estimate statistical significance of the top k shortest paths, and (4) Construct TopNet by retaining the edges in the significant shortest paths.

#### STEP 1, Integrate inputs

We integrate the inputs by computing (sample-specific or condition-specific) node and edge weights in the knowledge-based network using the omics data. In the specific scenario when comparing conditions (e.g. pre- and post-treatment), we encode the ‘response’ of the system to the change in conditions by assigning the node weight as either the fold change in gene expression (*N*_*i*_ = *FC*), or fold change in combination with simple gene expression (*N*_*i*_ = *SI* × *FC*). Here *N*_*i*_ is the weight of node *i*, and *SI* is the normalized signal intensity, or expression level, of a particular gene. Such a response can be in terms of up-regulated/activated pathways (Activated Response TopNet), obtained by computing *FC* = *SI*_*perturbed*_*/SI*_*control*_, or down-regulated/repressed pathways (Repressed Response TopNet), obtained using *FC* = *SI*_*control*_*/SI*_*perturbed*_. The Response TopNet is a union of these two TopNets, and provides a holistic view of the active, altered genes and processes. Exclusively applying the expression value as the node weight (*N*_*i*_ = *SI*) is useful either when no control is available, or when the emphasis is on identifying highly active processes in each state. This TopNet is referred to as the Highest Activity TopNet (HA TopNet). Even in this case, comparisons between states can be carried out after the TopNet is generated for each state.

To assign edge weight, we interpret an edge to represent a ‘reaction’ between the two nodes, and following the principles of mass action kinetics, an edge between highly abundant nodes is given Edge weight_(*i,j*)_ = *N*_*i*_ × *N*_*j*_, where *N*_*i*_ and *N*_*j*_ are the weights of the incident nodes *i, j*. This choice of edge weight prioritizes highly active interactions in a given context.

#### STEP 2, Compute top k shortest paths

To achieve a biological outcome, typically a sequence of active reactions is involved, represented by a series of high weight edges in our network. In order to enumerate such high weight paths, we first transform the edge weights into edge costs as Edge 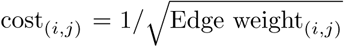, and use Dijkstra’s algorithm (Dijkstra *et al.* (1959)) to identify all-pair-shortest-paths. We then normalize the path cost for each node pair by the number of edges along the shortest path to get 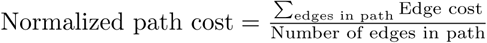, and retain the top *k* shortest paths, where *k* is a user-defined, application-specific threshold.

#### STEP 3, Statistical significance of shortest path costs

We assess the statistical significance of the normalized cost of each selected path empirically as follows. Given an *m* × *n* matrix of gene expression data for *m* genes in *n* samples/conditions, we randomly shuffle data in each row (gene) independently. The edges are re-weighted with the randomized gene expression data, and the cost of each path from step 2 is computed. After *r* such randomizations, for each path selected in step 2, *r* randomization-based costs are computed, based on which a z-score and p-value is estimated for each path. The p-value is finally transformed into a q-value (Benjamini and Hochberg (1995)) to account for multiple hypotheses testing. All paths with significant q-value are retained.

#### STEP 4, Construct TopNet

The edges in the significant paths from step 3 form a sub-network, which we refer to as the TopNet. The TopNet provides a snapshot of the active and/or significantly altered processes in the system, and can be studied to gain mechanistic insights. To further prioritize critical genes and paths in the TopNet, we apply network centrality measure—Ripple Centrality (Sambaturu *et al.* (2016)).

In cases where a single condition is being examined, or the number of conditions is too small to generate a sufficiently large number of randomized gene expression matrices, step 3 can be skipped, and top *k* shortest paths can be taken to represent highly active, altered paths, albeit without the statistical filter. In such cases, Step 4 can be directly applied to these paths to generate a TopNet.

### 2.2 Ripple centrality

Ripple centrality (Sambaturu *et al.* (2016)) prioritizes nodes which can reach a large fraction of the network along highly active and perturbed paths. It is measured as Ripple centrality(*u*) = *C*(*u*) × *R*_*out*_(*u*), where *R*_*out*_(*u*) = |nodes reachable from *u*| denotes the outward reachability of node *u*, and 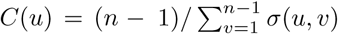 gives the closeness centrality of node *u*. Here *σ*(*u, v*) denotes the cost of the shortest path from node *u* to all *n* − 1 other nodes in the graph.

### 2.3 M.tb drug exposure

#### 2.3.1 Data

Transcriptomic data for M.tb H37Rv exposed for 16 hours to 2xMIC of 18 drugs was obtained from GSE71200 (Ma *et al.* (2015)). The list of 18 drugs along with their mechanism of action and TopNet details can be found in Supplementary Table S1. A knowledge-based network composed of experimentally validated protein-protein interactions as well as regulatory interactions in M.tb was obtained from (Mishra *et al.* (2017)), consisting of 3,686 genes and 34,223 edges.

#### 2.3.2 Gold standards

INH is known to affect the mycolic acid synthesis and processing pathways in M.tb (Wishart *et al.* (2017)). To create a gold standard for INH treatment, we searched for the term ‘mycolic acid’ in Mycobrowser (Kapopoulou et al. (2011)), a database of manually curated annotations for pathogenic mycobacteria, including M.tb. This resulted in a list of 17 M.tb genes, to which we added katG and fas, the known targets of INH (Wishart *et al.* (2017)). Similarly, gold standards were created for 5 other drugs by searching for terms related to their known mechanisms of action - ‘RNA polymerase’ for Rif, ‘mycolic acid’ for ethionamide, ‘protein synthesis’ for capreomycin, and ‘30s ribosomal protein’ as well as ‘16s rrna’ for kanamycin and streptomycin (Wishart *et al.* (2017)) (Supplementary Table S3).

#### 2.3.3 TopNet creation

For all 18 drugs in GSE71200 (Ma *et al.* (2015)), Activated Response TopNets were constructed using *N*_*i*_ = *SI*_*drug*_ × (*SI*_*drug*_*/SI*_*control*_), while *N*_*i*_ = *SI*_*control*_ × (*SI*_*control*_*/SI*_*drug*_) was used to construct the Repressed Response TopNets. Only shortest paths with 2 or more edges were considered, and 1,000 randomizations of the gene expression matrix were carried out for computing statistical significance of shortest paths. The percentile and q-value thresholds were chosen such that the resulting TopNets were of similar size for all cases (Supplementary Table S1). Activated and Repressed TopNets are provided in Supplementary Files S1 and S2, respectively.

#### 2.3.4 Functional enrichment

Functional enrichment was carried out using ClueGO v2.3.4 (Bindea *et al.* (2009)), a plugin in the network visualization tool Cytoscape 3.2 (Shannon *et al.* (2003)). Enrichment was against GO Biological Processes, GO Cellular Components and GO Molecular Functions, with a q-value cutoff of 0.01. Enriched pathways for all 18 drug exposure cases are provided in Supplementary File S3.

#### 2.3.5 Significance of TopNet connectedness

Significance of TopNet connectedness was tested by comparing against comparable sub-networks induced by (a) the top DEGs, (b) 1,000 sets of randomly sampled genes, and (c) 1,000 sets of randomly sampled edges. Here the number of DEGs and sampled genes (or edges) corresponds to the number of nodes (or edges) in the TopNet.

### 2.4 Human tissues

#### 2.4.1 Data

Normalized gene expression data was collected from GTEx (Carithers and Moore (2015)) (dbGaP accession number phs000424.v7.p2) for 39 human tissues, corresponding to 23 organs and 2 cell lines. The signal intensities of each tissue were summarized using the LMFit function in R (Limma package; Ritchie *et al.* (2015)). The antilog of the fitted value was used for further analysis as PathExt requires non-negative values. Human protein-protein interaction network (hPPIN) comprising regulatory, signaling and metabolic pathways was obtained from (Sambarey *et al.* (2017a)). This network has 17,062 proteins (nodes) and 208,759 interactions (edges).

#### 2.4.2 TopNet creation

Since no control was available, we constructed two types of TopNets - HA TopNets using *N*_*i*_ = *SI*, and z-score TopNets using *N*_*i*_ = |*z* − *score*| _*i*_. Here z-score for a gene *i* in a given tissue was computed with respect to all tissues, and statistical significance of shortest paths was computed by randomizing the |*z* − *score*| matrix 1,000 times. The size of the TopNet can vary across tissues and across percentile and false discovery rate thresholds. For the z-score TopNets, we explored percentile thresholds in the range [0.001, 1.0] and q-value thresholds from the set {0.001, 0.005, 0.01, 0.05} in each tissue to adjust the TopNet size to ≈ 300 nodes. Then for the HA TopNet of each tissue, we explored the same set of percentile thresholds so as to have a comparable size between HA and z-score TopNets; the percentiles across tissues were either 0.001 or 0.002 in all cases. Thresholds for all tissues are available in Supplementary Table S9. HA TopNets and z-score TopNets for all tissues are provided in Supplementary Files S4 and S5, respectively.

#### 2.4.3 Gold standards

The human protein atlas (HPA; Uhlén *et al.* (2015)), a compiled list of Disease genes (Feiglin *et al.* (2017)), and genes from the Disease Ontology browser of the Mouse Genome Informatics (MGI) database (Bult *et al.* (2018)) were used to validate the results. HPA provides lists of genes whose mRNA expression is elevated in a particular tissue. The elevated expression can correspond to one of three categories: (i) ≥ 5-fold mRNA levels in a particular tissue as compared to all other tissues, (ii) ≥ 5-fold higher mRNA levels in a group of 2-7 tissues, and (iii) ≥ 5-fold higher mRNA levels in a particular tissue as compared to average levels in all tissues. The union of genes from the above three categories form the gold standard. HPA data was downloaded on the 26^*th*^ of December, 2018. Disease genes were compiled by Feiglin *et al.* (2017) by cross-referencing data from two databases - Online Mendelian Inheritance in Man (OMIM, Hamosh *et al.* (2005)), and the Human Phenotype Ontology (HPO, Köhler *et al.* (2013)). OMIM is a compendium of associations between genetic variations and predominantly Mendelian disorders, while HPO provides a standardized vocabulary for working with such phenotypic abnormalities. The Disease Ontology browser of the MGI lists genes whose mutation is associated with phenotypes characteristic of human disease (Bult *et al.* (2018)). A list of housekeeping genes obtained from Eisenberg and Levanon (2013), comprising of 3,804 genes with constant expression level across a panel of tissues, is used as a negative control to test whether tissue TopNets are enriched in ubiquitously active genes.

#### 2.4.4 Functional enrichment and ranking of pathways

Enrichment was carried out using the enrichGO function of the R package clusterProfiler v3.6.0 (Yu *et al.* (2012)), using Biological Processes as the ontology, and with a Benjamini Hochberg cutoff of 0.01. For each tissue, the background for enrichment was set to be the list of genes for which both expression and interaction data were available. Pathway enrichment results for HA TopNets, z-score TopNets, their corresponding baselines, gold standards, as well as housekeeping genes, are provided for all tissues in Supplementary File S6. Pathways enriched in the TopNets were ranked based on the cost of the first TopNet shortest path involving a gene from that pathway. Ties were broken based on the fold enrichment of TopNet genes in a pathway relative to expectation.

## 3 Results

### 3.1 PathExt reveals pathways related to drugs’ mechanism of action in treated M.tb

In a previous study, the *Mycobacterium tuberculosis* (M.tb) strain H37Rv was exposed to different concentrations of various anti-tuberculosis drugs, and the transcriptional response was measured (GEO accession number GSE71200; Ma *et al.* (2015)). We obtained the transcriptomic data for 2xMIC (twice the minimum inhibitory concentration) dose of 18 drugs, for bacteria surviving 16 hours of drug exposure, suggesting a degree of drug resistance. Only one replicate per MIC per drug and a single untreated control sample were measured, making robust estimation of differential expression impractical. For 6 drugs where the mechanism of action is well studied (Wishart *et al.* (2017)), we obtained gold standard sets of genes experimentally verified to be perturbed upon drug exposure (Methods section 2.3.2). In all 6 cases, the Response TopNets generated by PathExt are concordant with the gold standards, and reveal genes and pathways relevant to the action of each drug (Table 1). In contrast, the genes with 1.5-fold differential expression have consistently poor overlap with gold standards (Table 1). We discuss the Isoniazid and Rifampicin exposures in detail below.

**Table 1:** Response TopNets for M.tb exposed to 6 drugs whose mechanism of action is well known, are concordant with gold standards and reveal genes relevant to the action of each drug.

| Drug | Gold standard | Activated Response TopNet |  |  | Repressed Response TopNet |  |  | Response TopNet |  |  | 1.5 FC DEGs |  |  |  |
| --- | --- | --- | --- | --- | --- | --- | --- | --- | --- | --- | --- | --- | --- | --- |
|  |  | Nodes | Gold standard | p-value | Nodes | Gold standard | p-value | Nodes | Gold standard | p-value | DEGs | Base network | Gold standard | p-value |
| Capreomycin | 29 | 184 | 2 | 0.45 | 195 | 5 | 0.02 | 372 | 7 | 0.03 | 1796 | 1676 | 16 | 0.26 |
| Ethionamide | 15 | 201 | 5 | 0.001 | 202 | 0 | 1 | 394 | 5 | 0.02 | 610 | 562 | 6 | 0.02 |
| Isoniazid | 17 | 195 | 6 | 0.0002 | 213 | 5 | 0.002 | 401 | 11 | 2.2e-7 | 1078 | 987 | 8 | 0.07 |
| Kanamycin | 30 | 191 | 5 | 0.02 | 199 | 2 | 0.5 | 379 | 7 | 0.03 | 790 | 731 | 4 | 0.23 |
| Rifampicin | 22 | 196 | 7 | 0.0001 | 197 | 4 | 0.03 | 380 | 11 | 4.3e-6 | 1579 | 1462 | 13 | 0.07 |
| Streptomycin | 30 | 193 | 7 | 0.0009 | 181 | 4 | 0.06 | 338 | 10 | 0.0003 | 308 | 286 | 1 | 0.29 |

#### 3.1.1 PathExt links INH exposure to mycolic acid synthesis and processing

The anti-bacterial drug Isoniazid (INH) inhibits the synthesis of mycolic acids, which are long fatty acids found in the cell walls of mycobacteria (Wishart *et al.* (2017)). The Activated Response TopNet (selecting for up-regulated paths), Repressed (down-regulated paths), and merged Response TopNets (Methods) identified by PathExt were all significantly enriched in gold standard genes related to mycolic acid synthesis and processing (Table 1, Supplementary Table S1). In stark contrast, the DEGs with ≥ 1.5-fold differential expression had poor overlap with the gold standard (Table 1, Supplementary Table S1). The central genes (Methods section 2.2) in the Activated Response TopNet consist mainly of genes involved in mycolic acid biosynthesis, whereas the Repressed Response TopNet has unsaturated acyl-CoA hydratases responsible for oxidizing fatty acids, and genes involved in lipid degradation as the central nodes (Supplementary Tables S4 and S5). These results unambiguously point to the up-regulation of fatty acid synthesis and down-regulation of its degradation as a resistance response to INH exposure.

A previous study (Takayama *et al.* (2005)) consolidated experimental and computational evidence to list the 7 main processes in the mycolic acid synthesis and processing pathway, namely, the FAS-I (fatty acid synthetase-I) system, transition from the FAS-I system to the FAS-II system, the FAS-II system, cyclopropane synthases and methyltransferases, oxidation-reduction, Claisen-type condensation, and mycolic acid processing. Of the 42 genes described in their work, interaction and expression data were available for 39, of which 16 were present in the INH exposed Response TopNet (3.63 fold enrichment; Fisher’s p-value = 1.68e-6), while the 1,078 DEGs comprise only 14 of these genes (Fisher’s p-value = 0.27). Notably, the TopNet sub-network induced by the 16 genes from the mycolic acid synthesis and processing pathway (Figure 2) and their immediate neighbors, represent all 7 component processes. Interestingly, NADH dehydrogenases (highlighted in violet in Figure 2) are also picked up in this sub-network. It has been hypothesized that M.tb may gain resistance to INH by regulating NADH dehydrogenase and the intracellular NADH/NAD+ ratio (Miesel *et al.* (1998)). This is consistent with the fact that the bacteria under study are the ones which survived exposure to 2xMIC of INH and thus likely to have triggered their resistance processes.

**Figure 2:**
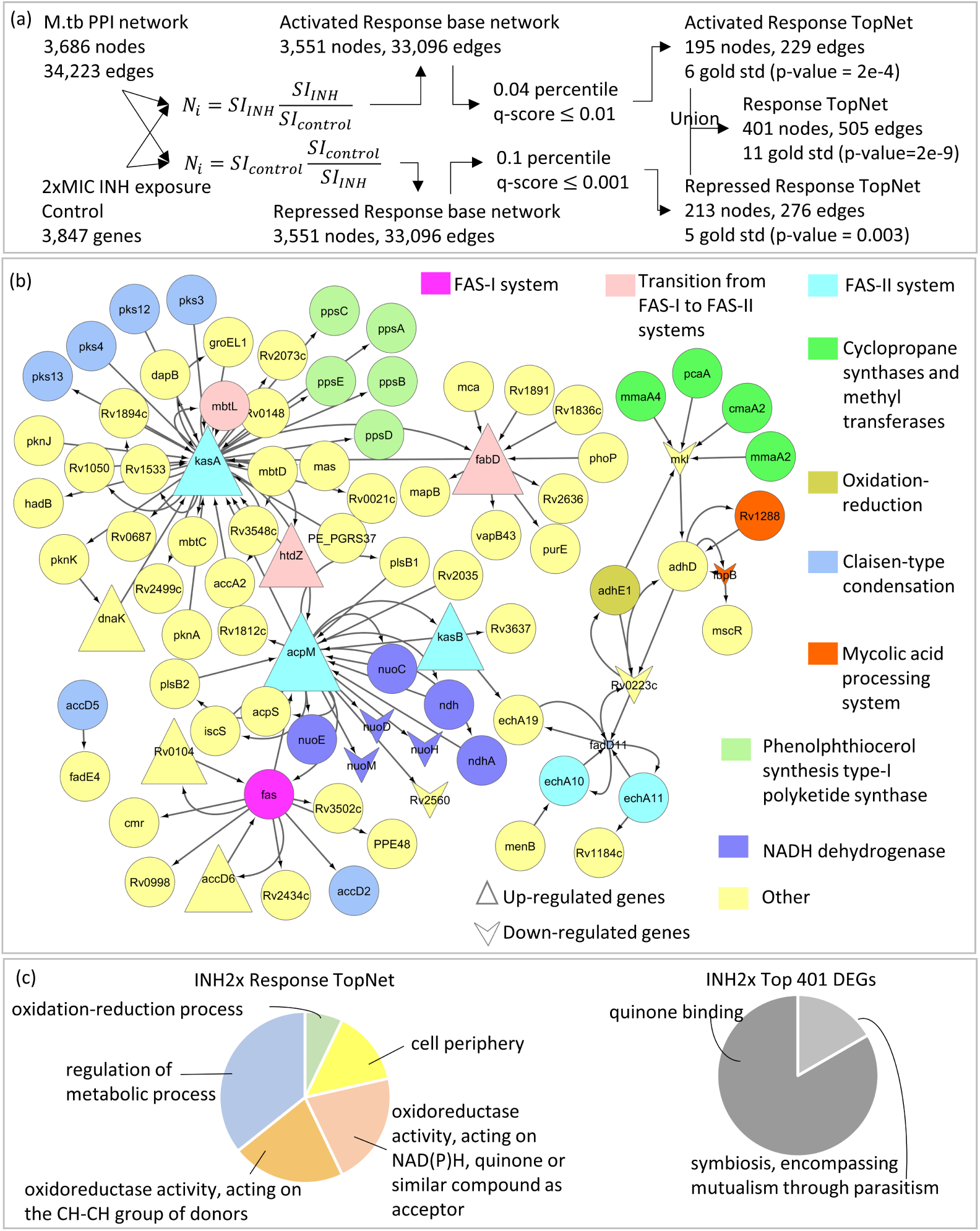
Response to 2xMIC INH. (a) Gene expression data for a single sample of M.tb exposed to 2xMIC of INH for 16 hours is integrated with a knowledge-based protein-protein interaction network for M.tb using two weighting schemes. *N*_*i*_ = *SI*_*I*_ *NH* × (*SI*_*INH*_ */SI*_*control*_) prioritizes genes up-regulated after exposure to INH, and results in an Activated Response TopNet. *N*_*i*_ = *SI*_*control*_ × (*SI*_*control*_*/SI*_*INH*_) prioritizes genes and processes down-regulated after exposure to INH, and results in a Repressed Response TopNet. Thresholds for the top *k* shortest paths and statistical significance are chosen such that the two TopNets are of comparable sizes. The union of the two TopNets gives a Response TopNet. All three TopNets are enriched in gold standard genes. (b) Sub-network of the Response TopNet formed by extracting genes from the mycolic acid synthesis and processing pathway (Takayama *et al.* (2005)), the known target pathway of INH, and their immediate interactors. Every component process of this pathway is represented in the Response TopNet by at least 1 gene. (c) GO enrichment of Response TopNet gives pathways relevant to INH exposure, such as cell-periphery and oxidation-reduction process. Enrichment of an equal number of top DEGs does not provide drug-specific insights.

Finally, as an additional control, we directly compared the Response TopNet genes with same number of top DEGs in terms of their functional enrichment (Figure 2, Methods section 2.3.4). The genes in the Response TopNet are enriched in the functional terms relevant to INH exposure, such as *cell periphery*, which is the part of the cell most affected by INH (Wishart *et al.* (2017)), and stress response terms such as *oxidoreductase activity* and *oxidation-reduction process*. We also find the term *regulation of metabolic processes*, which is an expected energy conservation response. In contrast, the top 401 DEGs are enriched for the terms *quinone binding* and *symbiosis encompassing mutualism through parasitism*, which are not informative of the condition under study. Together, these results show that the Response TopNet for M.tb exposure to 2xMIC of INH is indeed characteristic of its action and reveals genes and processes that would be missed by a conventional approach relying on differential gene expression alone.

#### 3.1.2 Rif exposure TopNet reveals the perturbation of nucleotide synthesis pathway

Rifampicin (Rif) inhibits DNA-dependent RNA polymerase activity, thus suppressing transcriptional initiation (Wishart *et al.* (2017)). Once again, the Activated, Repressed and union Response TopNets are enriched in gold standard genes, whereas the DEGs are not (Table 1, Supplementary Table S1). The gene rpoB (Rv0667) is central in the Activated Response TopNet, effectively recapitulating previous reports which suggest that Rif resistance can be caused by transcriptional up-regulation of rpoB (Zhu *et al.* (2018)). The error prone DNA repair synthesis protein DnaE2 (Rv3370c), and the genetic recombination and nucleotide excision repair protein RecA (Rv2737c) are also central in this network. Exposure to antibiotics such as Rif has been shown to result in a recA-dependent SOS response, and a corresponding increase in dnaE2 levels (McGrath *et al.* (2013)). Also, the up-regulation of dnaE2 has been identified as a critical factor in the emergence of drug resistance both *in-vitro* and *in-vivo* (Boshoff *et al.* (2003)). Other central genes (full list in Supplementary Table S4) include the 16S ribosomal RNA methyltransferase Rv2372c, and the replicative DNA helicase dnaB (Rv0058). These genes reflect perturbations in the nucleotide synthesis pathway, the very pathway known to be affected upon exposure to Rif. Central genes in the Repressed Response TopNet include, among others, dnaK (Rv0350) and Rv0232, a transcriptional regulator of the tetR/acrR-family. Disruption of Rv0232 has been shown to provide a growth advantage to H37Rv *in-vitro* (DeJesus *et al*. (2017)). We found that Rv0232 was 4.5-fold down-regulated and centrally involved in repressed paths, suggesting this as a possible resistance mechanism.

Interestingly, dnaK is central in the Repressed Response TopNet for Rif, whereas it is central in the Activated Response TopNet for INH exposure. It has been shown that dnaK is repressed by Rif (Eltringham *et al.* (1999)), whereas cells with higher levels of dnaK are more likely to persist upon exposure to INH (Jain *et al.* (2016)). This result underscores the biological and mechanistic relevance, as well as the condition-specificity of the TopNets generated by PathExt.

Although the exact pathway for DNA-dependent RNA polymerase activity is not known, examining the central genes from the Rif Activated and Repressed Response TopNets along with their immediate interactors provides valuable insights. These genes form two connected components, connected by two linker genes, fadE18 (Rv1933c) and fadD11 (Rv1550) (Figure 3). This sub-network highlights three major processes, namely, (i) transcription and nucleotide synthesis, (ii) error-prone synthesis and repair, and (iii) lipid metabolism. Figure 3 also shows the GO-term based enrichment of the genes in the Response TopNet, and for an equal number of top DEGs. The genes in the Response TopNet are enriched for terms relevant to exposure to Rifampicin, such as translation, which is the process targeted by Rif, plasma membrane and acyl-CoA dehydrogenase activity, which are related to lipid metabolism. On the other hand, the 380 top DEGs are enriched for the terms *cell periphery* and *plasma membrane*, which are not specifically informative of cellular response to the drug.

**Figure 3:**
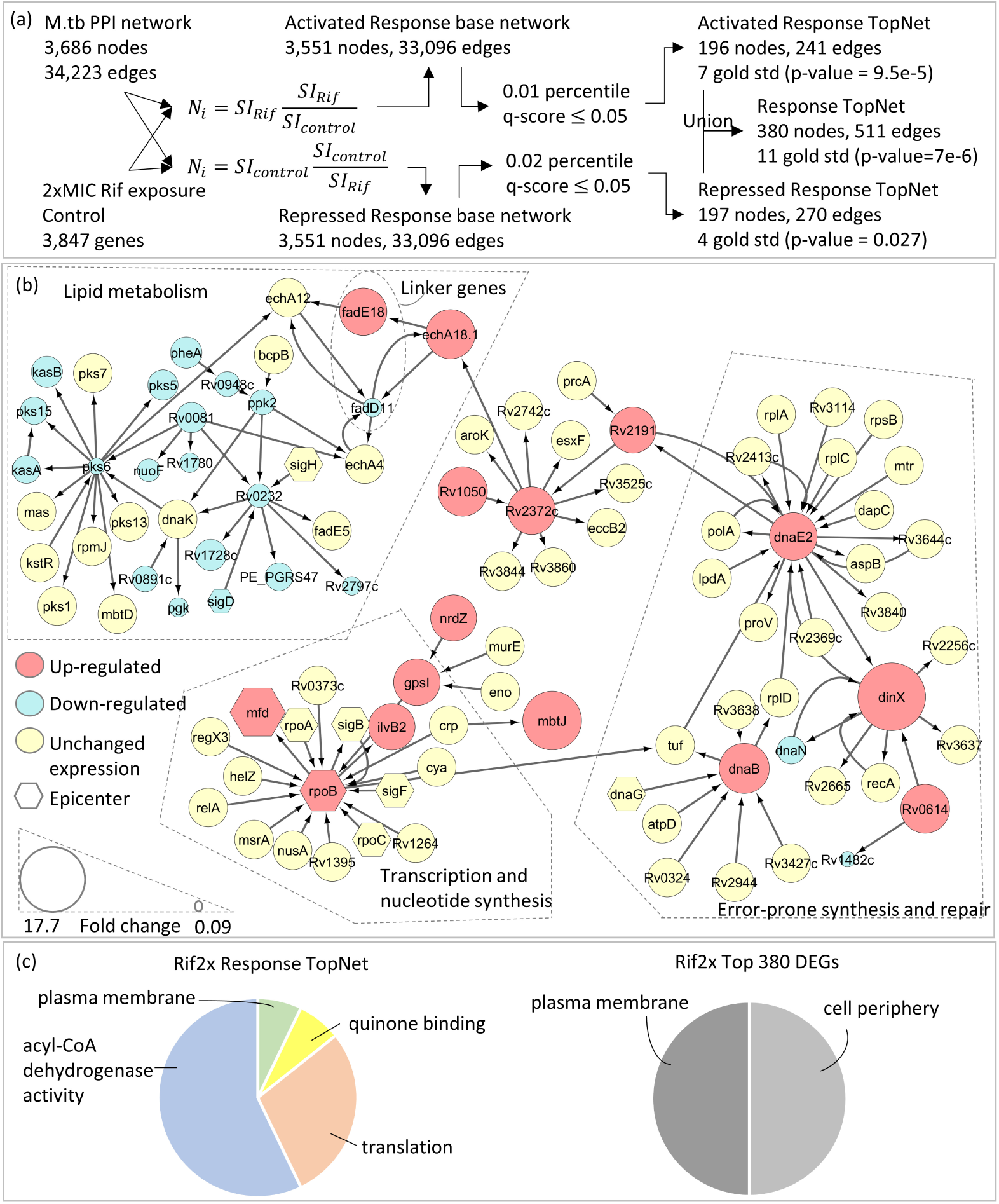
Response to 2xMIC Rif. (a) Gene expression data for a single sample of M.tb exposed to 2xMIC of Rif for 16 hours is integrated with a knowledge-based protein-protein interaction network for M.tb using two weighting schemes. *N*_*i*_ = *SI*_*Rif*_ × (*SI*_*Rif*_ */SI*_*control*_) prioritizes genes up-regulated after exposure to Rif, and results in an Activated Response TopNet. *N*_*i*_ = *SI*_*control*_ × (*SI*_*control*_*/SI*_*Rif*_) prioritizes genes and processes down-regulated after exposure to Rif, and results in a Repressed Response TopNet. Thresholds for the top *k* shortest paths and statistical significance are chosen such that the two TopNets are of comparable sizes. The union of the two TopNets gives a Response TopNet. All three TopNets are enriched in gold standard genes. Central genes from Activated and Repressed Response TopNets as well as their immediate interactors, extracted from the union Response TopNet for Rif. The inclusion of two linker genes, fadE18 and fadD11, links the resulting components into a single connected component. The size of the nodes reflects the extent of dysregulation of the genes. Up-regulated genes are colored red, while down-regulated genes are blue. This module contains genes related to transcription and nucleotide synthesis, the known target pathway of Rif (Wishart *et al.* (2017)). Other pathways represented here are lipid metabolism and error-prone synthesis and repair, both known mechanisms of resistance to Rif (Howard *et al.* (2018); Boshoff *et al.* (2003)). (c) GO enrichment of Response TopNet gives pathways relevant to Rif exposure, such as translation. Enrichment of an equal number of top DEGs does not provide drug-specific insights.

As demonstrated by the INH and Rif case studies, each Response TopNet reveals drug-specific mechanisms. Drug-specificity of the TopNets is further emphasized by the fact that there is no node or edge common to all 18 Response TopNets, despite the same knowledge-based network being used as input in all cases.

The Response TopNet is a connected graph with > 50% nodes in the largest component in each of the 18 drug exposures. This connectedness, reflective of biological pathways, is shown to be non-random (Methods section 2.3.5), and not captured by the sub-networks induced by top DEGs (Table 2, Supplementary Table S2). This suggests that our Response Network captures crosstalk between the dysregulated paths, which simple differential gene expression analysis may not.

**Table 2:**
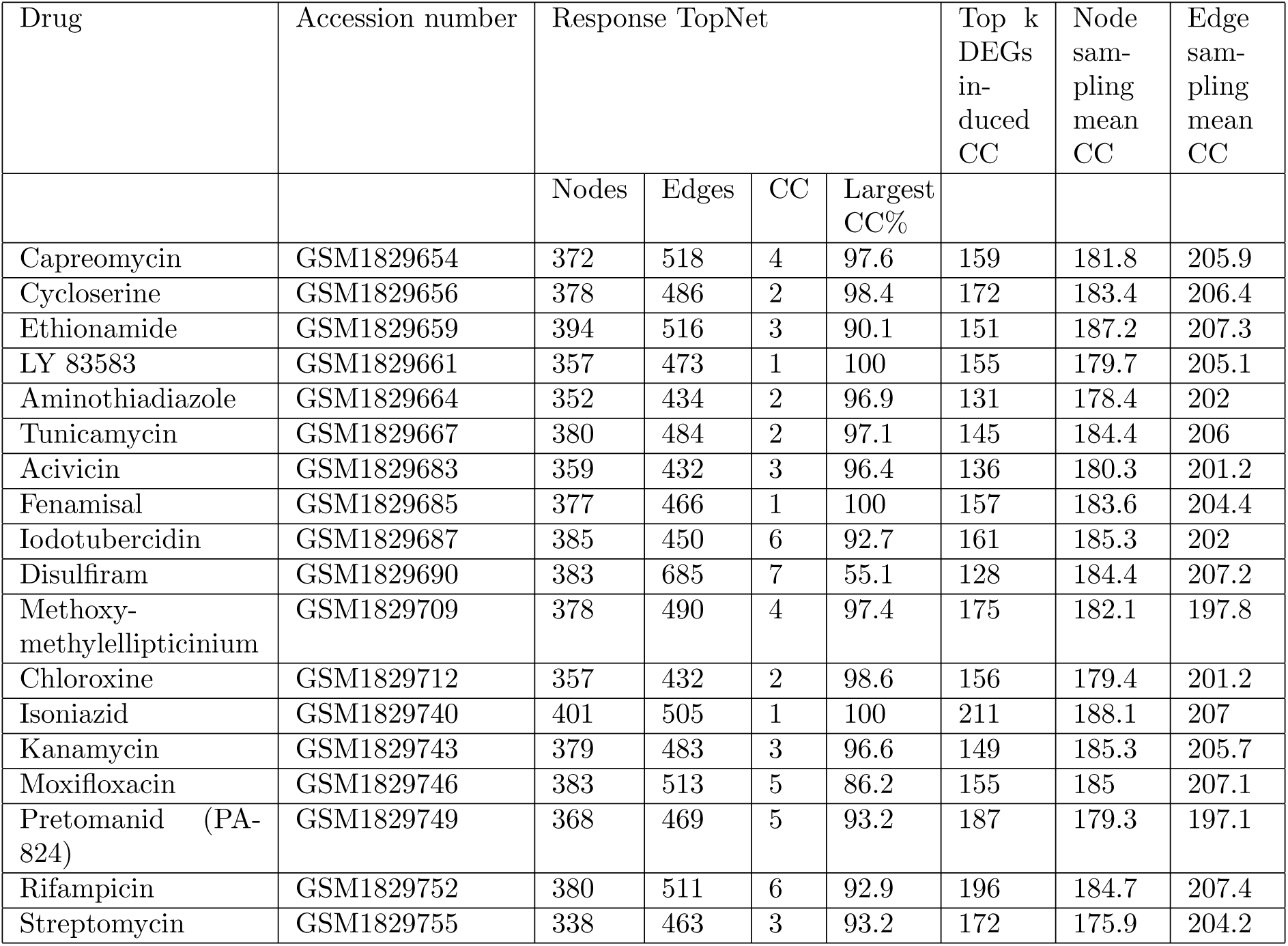
Response TopNets of all 18 drugs are connected graphs with > 50% nodes in a single connected component (CC). In contrast, the graphs induced by the same number of top DEGs, and by randomly sampling the same number of nodes or edges as the TopNet are highly disconnected. All randomizations are carried out 1,000 times.Standard deviations, z-scores and p-values are provided in Supplementary Table S2.

Taken together, these results show that PathExt captures drug-specific responsive genes and processes, even when only a single sample was available per condition.

### 3.2 Human tissue TopNets reveal tissue-related genes and processes

In a second set of case studies, we applied PathExt to identify tissue-related pathways using gene expression data for 39 human tissues in GTEx (Carithers and Moore (2015)), corresponding to 23 organs and 2 cell lines. In this scenario, there is no control. Therefore, we constructed two types of TopNets independently in each tissue (Methods section 2.4.2). A Highest Activity TopNet (HA TopNet) where *N*_*i*_ = *SI*, and a z-score TopNet where *N*_*i*_ = |*z* − *score*|_*i*_. Here *N*_*i*_ is the weight of node *i*, and *SI* is the normalized signal intensity (expression level). The z-score for gene *i* in a given tissue is computed relative to all tissues, thus using all tissues as a control for each tissue.

We assessed the tissue-specific TopNets against three gold-standards (Methods): (1) the Human Protein Atlas (HPA) (Uhlén *et al.* (2015)) where genes with ≥ 5-fold higher abundance in each tissue are labelled tissue-specific, (2) a set of curated tissue-relevant Disease genes (Feiglin *et al.* (2017)), and (3) a list of genes associated with tissue-specific human diseases from the MGI (Bult *et al.* (2018)). These comparisons are carried out for 37 out of 39 tissues, as corresponding gold standards could not be obtained for the 2 cell lines. We also use a list of housekeeping genes (Eisenberg and Levanon (2013)) as a negative control. To assess the utility of the z-score TopNets, we use the same number of genes with the highest |*z* − *score*| as a baseline control. Likewise, for the HA TopNets, the baseline used is the set of genes with highest expression levels.

The MGI had ≥ 25 genes with both gene expression and interaction data for 5 tissues. Of these, the HA TopNets were significantly enriched in tissue-associated genes in 3 tissues, and z-score TopNets in 4 tissues (Supplementary Table S6). In every case, the TopNet picked up equal or more gold standard genes than the corresponding baseline.

Figures 4 and 5 show the Fisher’s p-value of the overlap between the genes in the TopNets (and their corresponding baselines) and the two gold standards HPA and Disease genes. Since HPA is constructed based on differential abundance, as expected, genes with top z-score are highly concordant with the HPA-derived tissue-specific genes. In all other comparisons across tissues, genes identified by PathExt agree better with gold standards than the corresponding baselines. We found 4 exceptions out of 74 comparisons (37 tissues x 2 gold standards), marked with dashed boxes in Figures 4 and 5. Even in these cases, the pathways enriched in the TopNets are relevant to the functions of that tissue (Supplementary Info S1).

**Figure 4:**
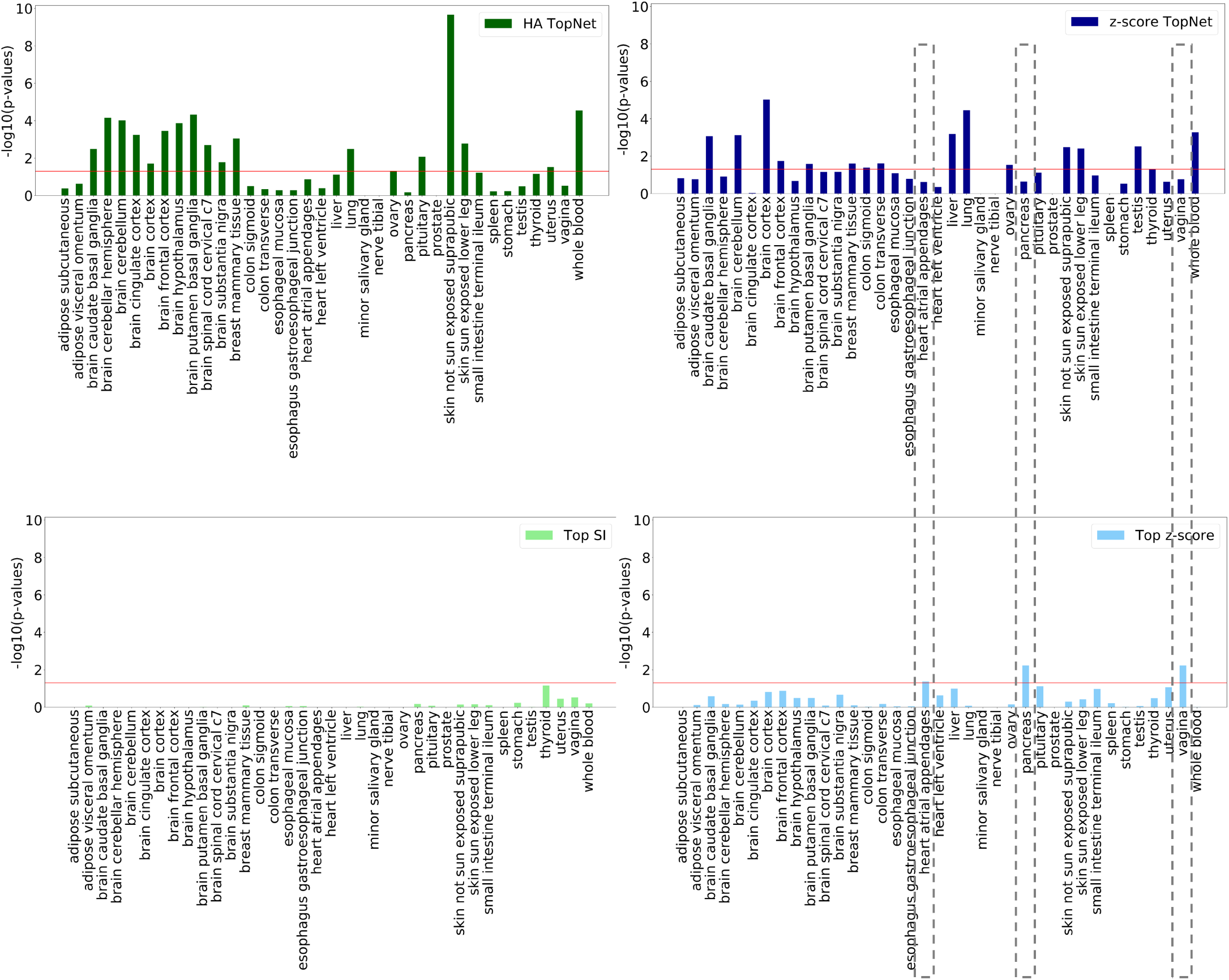
Overlap with Disease genes. p-values for overlap between the gold standard Disease genes, and nodes from (a) HA TopNets, (b) z-score TopNets, and their corresponding baseline controls, (c) Top SI (genes with highest expression) and (d) Top z-score. The p-values are plotted along the Y-axis as −*log*_10_(*p* − *value*), such that a taller bar corresponds to a more statistically significant overlap. The red horizontal line corresponds to p-value = 0.05. Tissues are plotted along the X-axis in lexicographic order. In every case except the three highlighted using dashed boxes, the overlap between TopNet nodes and the gold standard is better than the corresponding baseline. For pancreas and vagina, the genes with top z-score agree better with Disease genes than the genes in the z-score TopNet. These tissues are explored in greater detail in Supplementary Info S1.

**Figure 5:**
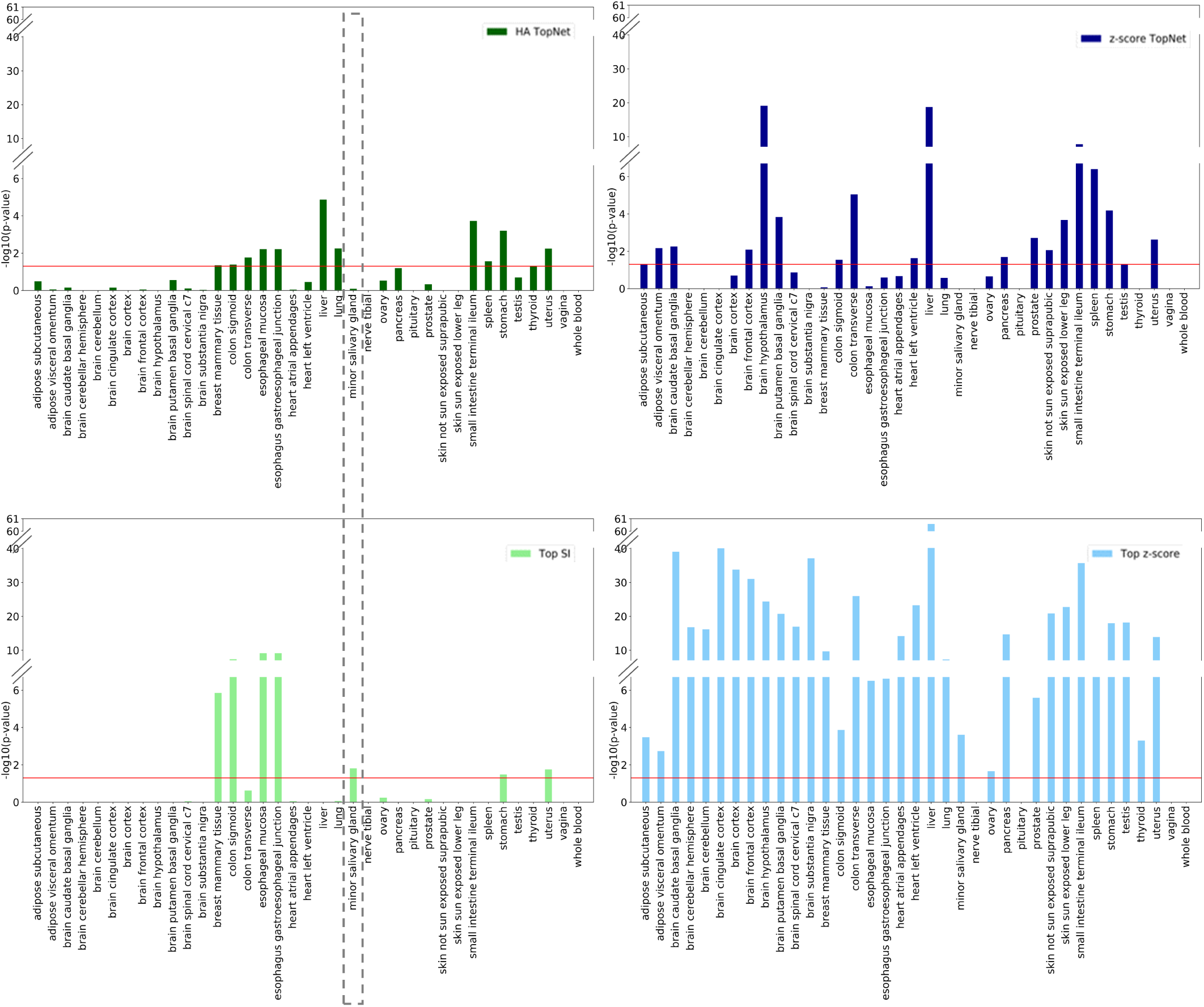
Overlap with HPA. p-values for overlap between the gold standard HPA and nodes from (a) HA TopNets, (b) z-score TopNets, and their corresponding baseline controls, (c) Top SI (genes with highest expression) and (d) Top z-score. The p-values are plotted along the Y-axis as −*log*_10_(*p* − *value*), such that a taller bar corresponds to a more statistically significant overlap. The red horizontal line corresponds to p-value = 0.05. Tissues are plotted along the X-axis in lexicographic order. Since genes with ≥ 5-fold differential expression are labelled tissue-specific in HPA, the genes with top z-score always have statistically significant overlap with this gold standard, as expected. Notwithstanding this comparison, the overlap between HA TopNet nodes and the gold standard is better than the corresponding baseline in all but one case (highlighted with dashed box); discussed in Supplementary Info S1.

An ideal tissue-specific network should exclude housekeeping genes, which by their very definition are broadly active. We find that the TopNets identified by PathExt have this property, and are under-enriched in housekeeping genes in all but 1 case (Supplementary Table S6). This suggests that the paths prioritized by PathExt correspond to tissue-related functions rather than universally active processes.

#### 3.2.1 PathExt-identified pathways enriched exclusively in a tissue correspond to known tissue-relevant functions

Figure 6 shows the top pathway exclusively enriched in the HA TopNet of each of 32 tissues (Methods section 2.4.4). The 7 excluded tissues had no exclusively enriched pathways. Figure 6 also highlights additional significant (0.01 ¡ q-value ≤ 0.05 and 0.05 ¡ q-value ≤ 0.1) pathway-tissue pairs. We found direct literature evidence supporting 20 out of 32 exclusive pathway-tissue associations, and indirect evidence for an additional 9 (Supplementary Table S7). A similar table along with literature evidence for the pathways enriched in the z-score TopNets is provided in Supplementary Table S8.

**Figure 6:**
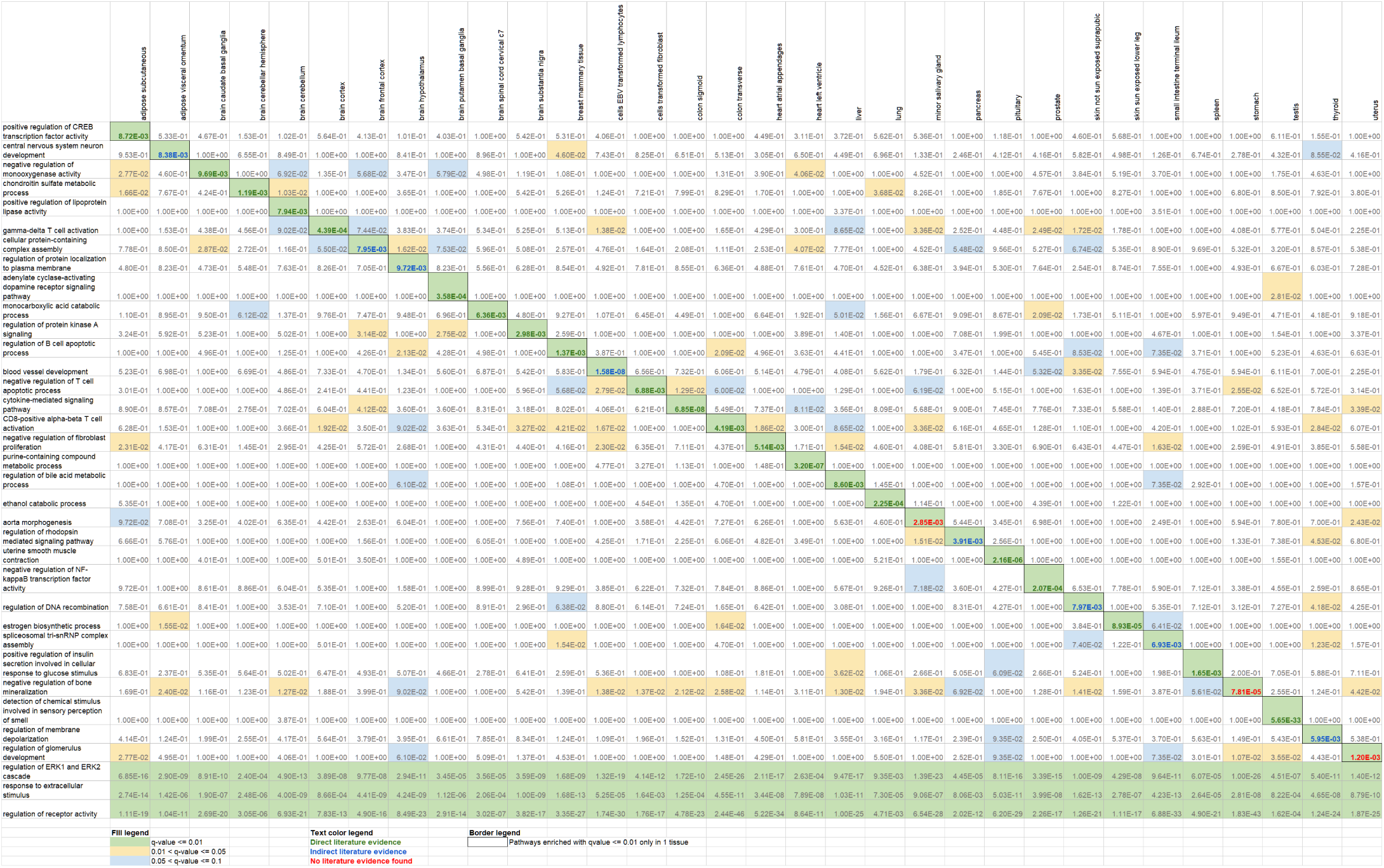
Tissue-exclusive pathways, HA TopNets. Top GO Biological Process exclusively enriched in each tissue, as well as the top 3 processes enriched in all tissues. The q-value of enrichment is provided for all cases. Green filling corresponds to cases with q-value ≤ 0.01, orange to 0.01 < q-value ≤ 0.05, and blue to 0.05 < q-value ≤ 0.1. Literature evidence support 29 out of 32 tissue-pathway pairs with q-value ≤ 0.01 (bold green text; Supplementary Table S7).

Some of the pathway-tissue pairs correspond to well-established functions of the tissue, such as *regulation of bile acid metabolic process* in liver (Chiang (2013)), *ethanol catabolic process* in lung (Bernstein (1982)), etc. PathExt reveals a few surprising associations as well. *Sensory perception of smell* is the top pathway exclusively enriched in the testis. At first glance this seems counter-intuitive. However, prevalence of olfactory receptors in the testis and sperm has been experimentally verified, and testicular olfactory receptor signaling has been implicated in sperm flagellar motility (Kang and Koo (2012)). As another example, *regulation of rhodopsin mediated signaling pathway* is enriched exclusively in the pancreas. Interestingly, rhodopsin regulates insulin receptor signaling in rod photoreceptor neurons (Rajala and Anderson (2010)), and loss of Arf4, a GTPase important for localizing rhodopsin to the eye and kidney, has been shown to result in damage of exocrine pancreas in mice (Pearring *et al.* (2017)). This surprising link between rhodopsin and the pancreas is not picked up by any of the gold standards or the controls.

Figure 6 also highlights the specificity of functions of the different regions of the brain. For instance, the *gamma-delta T cell activation pathway* is enriched in the brain cortex. Gamma-delta T cells have been implicated in Rasmussen encephalitis, a disease characterizing inflammation of the cerebral cortex (Owens *et al.* (2015); Varadkar *et al.* (2014)). The *adenylate cyclase-activating dopamine receptor signaling pathway* is enriched exclusively in the putamen basal ganglia region of the brain. The dorsal region of the basal ganglia comprises of the putamen, and the caudate nucleus (Lanciego *et al.* (2012)). Experiments involving homogenates of the caudate nucleus of the rat brain point at dopamine-sensitive adenylate cyclase as the receptor for dopamine in the mammalian brain (Kebabian *et al.* (1972)). This finding could indicate the presence of caudate nucleus cells in the putamen sample, or a shared function between these two adjacent regions of the brain. Several processes expected to be ubiquitous, such as *regulation of receptor activity* and *response to extracellular stimulus*, are enriched in all the tissues under consideration.

Overall, PathExt-identified tissue-specific TopNets recapitulate gold standard genes with known tissue-specific functions, and provide unique insights into tissue functions, not reflected in conventional differential expression-based analyses.

## 4 Discussion

We provide PathExt, a computational tool to identify sub-networks of an omics-integrated biological network, which capture the response to a perturbation, or the active processes in a particular condition. PathExt builds on our prior work which mined omics-integrated networks to (i) identify tuberculosis biomarkers (Sambarey *et al.* (2017b)), (ii) discriminate between primary and metastatic melanoma (Metri *et al.* (2017)), and (iii) identify influential genes in the condition under study (Sambaturu *et al.* (2016)). Substantially extending our prior work, PathExt provides a general framework to address all the above questions, while employing rigorous statistical significance estimation to identify critical paths. Importantly, PathExt is designed to operate even when a single sample is available for each condition, and in the absence of an experimental control sample.

Current approaches to identify active sub-networks are largely built on the work by Ideker *et al.* (2002), called jActiveModules, which formulates a sub-network scoring scheme based on the statistical significance of differential gene expression, and then identifies high-scoring sub-networks using a simulated annealing approach. Cabusora *et al.* (2005) use the same scoring method but identify sub-networks by listing k-shortest paths in the interaction network among a set of ‘seed’ nodes. The best scoring sub-network is then identified by sampling seed nodes based on their differential expression. Although this method computes paths (unlike jActiveModules), all edges in the interaction network are given equal importance, and the scoring scheme as well as seed node prioritization still focuses on DEGs. Other methods along similar ideas have been proposed, that filter sub-networks based, for example, on network motifs (Milo *et al.* (2002)). In contrast, PathExt assigns weights to the interactions in the biological network as a function of the given omics data, thus transferring importance from individual genes to paths, and potentially capturing the way in which biological phenotypes emerge from interconnected processes. Interestingly, even though connectedness is not used as a criterion to identify sub-networks, the TopNet resulting from the identified paths forms a well-connected graph.

PathExt relies on two user defined parameters, the threshold *k* used to select the top *k* shortest paths, and the q-value for statistical significance of the paths selected to construct TopNet. These values have been set at very stringent values in this paper, allowing us to focus on the most active paths. Different thresholds can give different layers of information, with different levels of false discovery.

In summary, PathExt is a general framework for path-based mining of omics-integrated biological networks. While the paths identified by PathExt may not constitute a comprehensive or exhaustive listing of all the active, altered processes in the system, the resulting TopNet can be thought of as a starting point from which hypotheses can be generated. In this work, we have gathered, for each drug and each tissue, the top central genes, along with their fold change for drug exposure (Supplementary Tables S4, S5), and z-score for human tissues (Supplementary Tables S10, S11). Further examining the network or genomic neighborhood of these and other genes comprising the TopNet can provide additional insights, or strengthen the insights gained.

## Funding

This work was supported by the Department of Biotechnology (DBT) - Indian Institute of Science (IISc) Partnership Program - Phase II [BT/PR27952/IN/22/212/2018] and the Mathematical Biology Initiative [DSTO/PAM/GR-1303] of the Government of India. S.H. was supported in part by National Science Foundation (NSF) award 1564785 and in part by the Intramural Research Program of the National Cancer Institute, Center for Cancer Research, National Institutes of Health (NIH).

## Notes

#### Summary of Updates

Author order updated

