## Supplementary material for "PathExt: a general framework for path-based mining of omics-integrated biological networks": Supplementary_Info_S1.docx

We scrutinized the four tissues (pancreas, minor salivary gland, heart atrial appendages, and vagina) where the TopNet had a poorer overlap with gold standards than their corresponding baselines (Fig. 5,6). The top three pathways enriched in each of these tissues, and also in the uterus, are shown in Table 1. Uterus is included for comparison with the TopNet of vagina.

Pancreas: Enrichment of Arginine methylation in the z-score TopNet of the pancreas is supported by the identification of arginine methyltransferase 1 (PRMT1) as a potential drug target for pancreatic cancer (Gao *et al.* (2019)). The HA TopNet is enriched in multiple pathways related to growth hormone signaling, which is a well-established process in pancreatic beta cells, impacting insulin release (Kawabe and Morgan (1983)). In contrast, pancreas-specific genes reported in HPA and Disease genes are enriched in digestion, lipid digestion, or gland development, which are not specific to the pancreas.

Minor salivary gland: In HPA, salivary gland (minor salivary gland is not represented in HPA), is enriched in expected pathways such as saliva secretion and sensory perception of taste. Even though neither of the TopNets is enriched in these pathways (this may reflect difference between major and minor salivary glands), interestingly, they are uniquely enriched in ethanol metabolism (z-score TopNet), and immune response (HA TopNet). It has been shown that exposure to alcohol-derived acetaldehyde, the first metabolite of ethanol, occurs in the oral cavity independently of metabolism in the liver, and is an independent risk factor for cancer (Stornetta *et al.* (2018)). The salivary gland is also known to be an active site for immune response (Deslauriers *et al.* (1986)) and has even been considered as an immunization site (Ponzio and Sanders (2017)).

Heart atrial appendages: Atrial appendages of the heart are tiny sacs projecting out of the atrial walls, and are distinct from the atrium proper in structural and physiological properties (Al-Saady *et al.* (1999)). Thus, it is to be expected that the TopNets corresponding to the atrial appendages will differ from the gold std genes specific to the heart itself. Although the genes with top |z-score| overlap well with the Disease genes related to the heart (p-value = 0.04), the top 3 pathways enriched for these top DEGs are regulation of receptor activity, chemokine-mediated signaling pathway, and neutrophil migration, which are not specific to the heart or its appendages. On the other hand, the pathways enriched by the z-score TopNet are related to glutathione derivative metabolism. This is supported by the fact that glutathione-sensitive attenuation of calcium current has been observed in monocytes derived from atrial appendages in patients with atrial fibrillation (Carnes *et al.* (2007)).

Vagina: We find that the TopNets of the uterus and the vagina are very similar to each other (>80% common nodes in z-score TopNets, and >70% common nodes in HA TopNets). Moreover, the enriched pathways, such as regulation of renal sodium excretion, regulation of homeostasis, and blood coagulation reflect functions typically associated with the urogenital system (Table 1, Paz and West (2013)). Similar enrichment is observed in the genes specific to uterus as per the gold standard Disease genes (Table 1).

Table 1. Top 3 enriched pathways in 4 select tissues. The tissues are pancreas and vagina, for which the genes with top z-score compared better with gold std Disease genes than their z-score TopNets, and minor salivary gland, for which the genes with top expression compared better with gold std HPA than the HA TopNet. Uterus is included for comparison with the pathways enriched in the tissue vagina.

| Pancreas | |
| --- | --- |
| z-score TopNet | peptidyl-arginine N-methylation; histone arginine methylation; ethanol oxidation |
| Top z-score | digestion; epithelial cell differentiation; regulation of hormone levels |
| HA TopNet | JAK-STAT cascade involved in growth hormone signaling pathway; growth hormone receptor signaling pathway; cellular response to growth hormone stimulus |
| Top Expression | regulation of receptor activity; leukocyte migration; monocyte chemotaxis |
| HPA | digestion; lipid digestion |
| Disease genes | gland development; negative regulation of epithelial cell proliferation; digestive system development |
| Minor salivary gland | |
| z-score TopNet | ethanol oxidation; ethanol metabolic process; regulation of transcription from RNA polymerase II promoter involved in heart development |
| Top z-score | chemokine-mediated signaling pathway; xenobiotic metabolic process; cornification |
| HA TopNet | chemokine-mediated signaling pathway; natural killer cell chemotaxis; lymphocyte chemotaxis |
| Top Expression | antimicrobial humoral response; regulation of receptor activity; neuropeptide signaling pathway |
| HPA | saliva secretion; secretion by tissue; detection of chemical stimulus involved in sensory perception of taste |
| Heart atrial appendages |  |
| z-score TopNet | glutathione derivative metabolic process; glutathione derivative biosynthetic process; ethanol oxidation |
| Top z-score | regulation of receptor activity; chemokine-mediated signaling pathway; neutrophil migration |
| HA TopNet | chemokine-mediated signaling pathway; estrous cycle; natural killer cell chemotaxis |
| Top Expression | regulation of receptor activity; feeding behavior; antimicrobial humoral response |
| HPA | muscle contraction; muscle system process; striated muscle contraction |
| Disease genes | heart development; cardiac muscle contraction; muscle contraction |
| Vagina | |
| z-score TopNet | positive regulation of renal sodium excretion; renal sodium excretion; regulation of renal sodium excretion |
| Top z-score | anterior/posterior pattern specification; regionalization; pattern specification process |
| HA TopNet | blood coagulation, fibrin clot formation; negative regulation of blood coagulation; negative regulation of hemostasis |
| Top Expression | regulation of receptor activity; antimicrobial humoral response; humoral immune response |
| Disease genes | embryonic morphogenesis; tube morphogenesis; tube development |
| Uterus | |
| z-score TopNet | G-protein coupled acetylcholine receptor signaling pathway; blood coagulation; intrinsic pathway, phasic smooth muscle contraction |
| Top z-score | anterior/posterior pattern specification; regionalization; embryonic skeletal system development |
| HA TopNet | blood coagulation, intrinsic pathway; blood coagulation, fibrin clot formation; neuropeptide signaling pathway |
| Top Expression | regulation of receptor activity; digestion; antimicrobial humoral response |
| HPA | embryonic limb morphogenesis; embryonic appendage morphogenesis; appendage morphogenesis |
| Disease genes | mismatch repair; urogenital system development; developmental growth |
